# Depolarizing afterpotentials in rodent mEC layer II neurons vary gradually across all principal cell types

**DOI:** 10.1101/2025.02.13.638108

**Authors:** Franziska S. Kümpfbeck, Johannes Nagele, Felix Felmy, Andreas V.M. Herz

## Abstract

Principal cells in Layer II (LII) of the rodent medial entorhinal cortex (mEC) often discharge in short burst episodes, whose biophysical mechanism and biological function are not fully understood. As shown by *in vitro* recordings in rats and *in vivo* recordings in mice, action potentials of stellate cells (SCs) and pyramidal cells (PCs) in mEC LII can trigger depolarizing afterpotentials (DAPs) that facilitate spike doublets and burst firing. *In vitro* recordings in mice confirm these findings for SCs and two additional cell types intermediate between SCs and PCs but suggest that mouse PCs generate minute DAPs only. To better understand the diversity and physiology of the intermediate LII mEC cell types, we performed slice experiments in Long Evans rats of both sexes and analyzed the data using clustering techniques. The analysis suggests four distinct cell clusters, reminiscent of previous observations in mice. On purpose, DAP properties were not used for the cluster analysis so that they could be studied post-hoc in an unbiased manner. Our results show that characteristic features of DAPs and other physiological markers are broadly distributed within each cell class and vary gradually from cluster to cluster, forming a continuum of biophysical properties.

## Introduction

The information processing of single neurons and larger neural networks is influenced by cell-type-specific membrane-potential dynamics. Sag potentials enable neurons to respond with short latencies to stimulus onsets, to exhibit subthreshold membrane potential oscillations (MPOs), and to resonate with oscillatory current inputs (Alonso and Klink, 1993; Puil et al., 1986; Erchova et al., 2004). Depolarizing afterpotentials (DAPs) create windows of opportunity for repeated firing that result in doublets of action potentials (APs) or brief AP bursts (Alonso and Klink, 1993; Canto and Witter, 2012; Csordás et al., 2020). These, in turn, play an important role in brain function, for example by boosting transmitter release that facilitates long-term potentiation in entorhinal-hippocampal circuits (Nicoll and Schmitz, 2005).

Traditionally, layer II (LII) of the rodent medial entorhinal cortex (mEC) has been assumed to consist of two strictly separated principal cell types, stellate cells (SCs) and pyramidal cells (PCs). Both mEC cell types were first characterized by Alonso and Klink (1993) using *in vitro* recordings in rats. Mediated by hyperpolarization-activated currents (*I*_*h*_), SCs display a prominent sag potential in response to a current step, often resulting in rebound spikes or bursts (Alonso and Klink, 1993). In addition, these currents support subthreshold MPOs and electrical resonances at frequencies in the theta range which may play a role in frequency-dependent information flow (Alonso and Klink, 1993; Erchova et al., 2004, Canto and Witter, 2012; Alessi et al., 2016). A large fraction of SCs show pronounced DAPs and AP doublets as well as short bursts whose inter-spike intervals reflect the time lag between an AP and the DAP it triggers (Canto and Witter, 2012; Csordás et al., 2020).

In contrast, PCs from LII mEC display neither membrane oscillations nor strong resonance properties. PCs that show no sag in response to hyperpolarizing currents also do not show a sag in response to depolarizing currents. These PC cells have small rebounds and large AP latencies following depolarizing step-current inputs, unlike SCs, which fire with short latencies to such inputs. Moreover, PCs differ from SCs by their pyramidally shaped soma with a single apical dendrite (Canto and Witter, 2012).

Although rat SCs and mouse SCs show rather similar characteristics, some PC properties seem to differ markedly in both species. In rats, PCs tend to exhibit substantial sag potentials (Canto and Witter, 2012) whereas this does not seem to be the case in mouse PCs (Fuchs et al., 2016), but see Winterer et al. (2017), whose data indicate roughly equal sag potentials for mouse SCs and PCs. In addition, the majority of rat PCs elicits DAPs of more than 2 mV (Canto and Witter, 2012; Alessi et al., 2016), while the reported DAP amplitudes in mice are much smaller, ranging between 0.04 ± 0.05 mV (mean ± SD) as reported by Fuchs et al. (2016) and 0.26 ± 0.44 mV (mean ± SD) as reported by Winterer et al. (2017). Finally, Ferrante et al. (2017) do not observe DAPs in mouse PCs and only in roughly 15% of mouse SCs.

Intermediate cell classes in between SCs and PCs have been found, too. Based on morphological features, Canto and Witter (2012) suggested the existence of three such classes in rat mEC Layer II, and Fuchs et al. (2016) proposed two groups for mice, named intermediate SCs (IMSCs) and intermediate PCs (IMPCs). To come to this conclusion, Fuchs et al. performed a principal component analysis (PCA) that took into account the following five electrophysiological and morphological features: (1) hyperpolarizing and depolarizing sag potentials, (2) burst firing in response to a step depolarization, as measured by the ratio of the first to second inter-spike interval (ISI1/ISI2 ratio), (3) depolarizing afterpotentials (DAPs), (4) latency of the first AP upon step depolarization and (5) the presence of a main (apical) dendrite. The first three criteria had also been used previously to differentiate between stellate and non-stellate mEC cells (Alonso and Klink, 1993; Canto and Witter, 2012).

To further categorize EC LII neurons, Varga et al. (2010) identified two cellular markers, Calbindin and Reelin, which label PCs and SCs, respectively. In mice, IMSCs express Reelin with a minority co-expressing Calbindin. Similarly, different varieties of IMPCs have been found: Calbindin positive, Reelin positive, and Reelin as well as Calbindin positive (Fuchs et al., 2016). IMPCs have pyramidally shaped somata, oriented horizontally with one main spiny apical dendrite. IMSCs differ from SCs in that they display a longer latency of the first AP upon depolarization near threshold (> 100ms). According to Fuchs et al. (2006), IMPCs exhibit pronounced DAPs (1.56 ± 0.22 mV; mean ± SD) roughly forty times the average amplitude of DAPs in PCs.

The discovery of intermediate cell classes demonstrates that principal neurons in rodent mEC LII come in more flavors than originally assumed. In addition, comparing the findings of Canto and Witter (2012) with those of Fuchs et al. (2016) and Winterer et al. (2017) shows that a number of open questions remain: What are the fundamental differences between rat and mice principle cells regarding sag potentials and depolarizing afterpotentials? How many cell classes would an in-depth analysis suggest? Are these classes compatible with the four (Fuchs et al., 2016) or five (Canto and Witter, 2012) groups reported for mice and rats, respectively? Or are there just two groups, as originally proposed by Alonso and Klink (1993) and again suggested by Winterer et al. (2017)? Finally, how would all the results fit with *in vivo* data (Domnisoru et al., 2013; Latuske et al., 2015; Csordás et al., 2020)?

## Materials and Methods

### Animals

Experiments were conducted with 80 Long-Evans rats (Charles River Laboratories International, Inc.) of both sexes that were housed in separate rooms together with their siblings in 612 × 435 × 216 mm boxes (internal floor area: 2065 cm^2^, Typ 2000P), filled with wooden chipping and wooden wool as bedding. Animals were kept under constant conditions with a 12 hour light/dark cycle, a room temperature of 22 - 23 ^°^C and a humidity of 50 - 55%. Animals were allowed to eat ad libitum and had unrestricted water access. All experiments complied with the institutional guidelines, regional and national laws.

### Preparation

Sagittal and horizontal brain slices containing the mEC were prepared from postnatal day P30-P45 animals. They were anesthetized with isoflurane and rapidly decapitated. Brains were quickly removed and transferred to ACSF containing (in mM) 50 or 120 sucrose, 25 NaCl, 27 NaHCO3, 2.5 KCl, 1.25 NaH2 PO4, 3 MgCl2, 0.1 CaCl2, 25 glucose, 0.4 ascorbic acid, 3 myo-inositol and 2 Na-pyruvat (pH 7.4 when bubbled with 95% O2 and 5% CO2). Brain slices of 400 *μ*m were taken using a VT1200S vibratome (Leica Microsystems GmbH, Wetzlar, Germany) and incubated in recording solution (same as ACSF but with 125 mM NaCl, no sucrose, 1.2 mM CaCl2, and 1 mM MgCl2) at 36^°^C for 45 minutes.

### Electrophysiology

After incubation slices were transferred to a recording chamber and continuously perfused with recording solution. Whole-cell recordings were acquired from 531 visually identified mEC neurons of layer II using a CCD camera (TILL-Imago VGA). A TILL Photonics imaging system (Gräfelfing, Germany), a combination of a charge-coupled device camera and a monochromator (Polychrome IV), was mounted onto an upright microscope (BX50WI) with a 60 × 1 numerical aperture (NA) objective (Olympus, Center Valley, PA). The microscope was equipped with gradient contrast illumination (Luigs and Neumann, Ratingen, Germany) and a 1.4-NA oil-immersion condenser. Recordings were performed with an EPC10/2 amplifier (HEKA Elektronik, Lambrecht/Pfalz, Germany).

All experiments were performed at physiological temperature of 36 ± 1^°^C, which was maintained by bath chamber (PH-1) and in-line (SF-28) heaters (Warner Instruments, Biomedical Instruments) and controlled by a temperature sensor placed next to the slice. All recordings were performed with 3-3.5 MΩ filamented, fire-polished borosilicate glass electrodes (inner diameter: 0.86 mm; outer diameter: 1.50 mm, Harvard Apparatus Ltd., Edenbridge, Kent, UK), pulled with a horizontal multi-step puller (DMZ-Universal Puller, Zeitz Instruments, Martinsried, Germany) and heat-polished to guarantee a smooth tip. Electrodes were filled with internal solution and fixated to an electrode-holder/head-stage, which was connected to a PatchStar Micromanipulator (ScientificaLtd., Uckfield, East Sussex, UK) to translate xyz direction, and to a manometer to control orally-generated pipette pressure. This positive pressure was applied to retain a clean pipette tip. The ground electrode and the electrode linking the head-stage with the internal pipette solution were made of Ag/AgCl wire. For current-clamp recordings, an internal solution containing (in mM) 145 K-gluconate, 4.5 KCl, 15 HEPES, 2 Mg-ATP, 2 K-ATP, 0.3 Na-GTP, 7.5 Na2-phosphocreatin adjusted by adding KOH to pH 7.25 was used.

To reconstruct and verify the location of recordings, cells were filled with 100 *μ*M Alexa 594 added to the internal solution. During current-clamp recordings the bridge balance was compensated to 100% after the series resistance had been estimated. The liquid junction potential (LJP = 17 mV, see Ammer et al., 2015) was not corrected for any of the solutions. Data were acquired at 20-25 kHz and filtered at 3 kHz. At the end of the recording the pipette was slowly and carefully retracted until a sufficiently high resistance was reached to ensure the cell membrane re-sealed and to avoid damaging the cell. Finally slices were fixated in 4% PFA (paraformaldehyde) for 3 hours.

### Stimulus generation

Stimulus generation and data acquisition were executed primarily with the Patchmaster software (HEKA Elektronik, Lambrecht (Pfalz), Germany). The stimulus template for the ZAP and sine-wave stimuli was generated with two other programs (*Template Creator*, costum C^++^ software by C. Kirst, LMU Munich, Germany and *IgorPro*, from WaveMetrics, Lake Oswego, OR).

#### Membrane resistance and time constant

To determine the membrane resistance and time constant, a 400 ms hyperpolarizing current step of -5 pA was injected at the cell’s resting membrane potential and repeated 60 times. The membrane resistance *R* was calculated by dividing the time- and trial-averaged voltage deflection by the applied current. The trial-averaged voltage response was fitted in the time interval from stimulation onset to peak hyperpolarization by an exponential function whose time constant was identified with the membrane time constant *τ*.

#### Sag potential

Sag potentials were measured in response to a hyperpolarizing current injection of -150 pA. The sag was calculated as the voltage difference between the membrane potential at steady state (last 200 ms) and the maximum membrane potential hyperpolarization (Fig. 1*A*).

**Figure 1.**
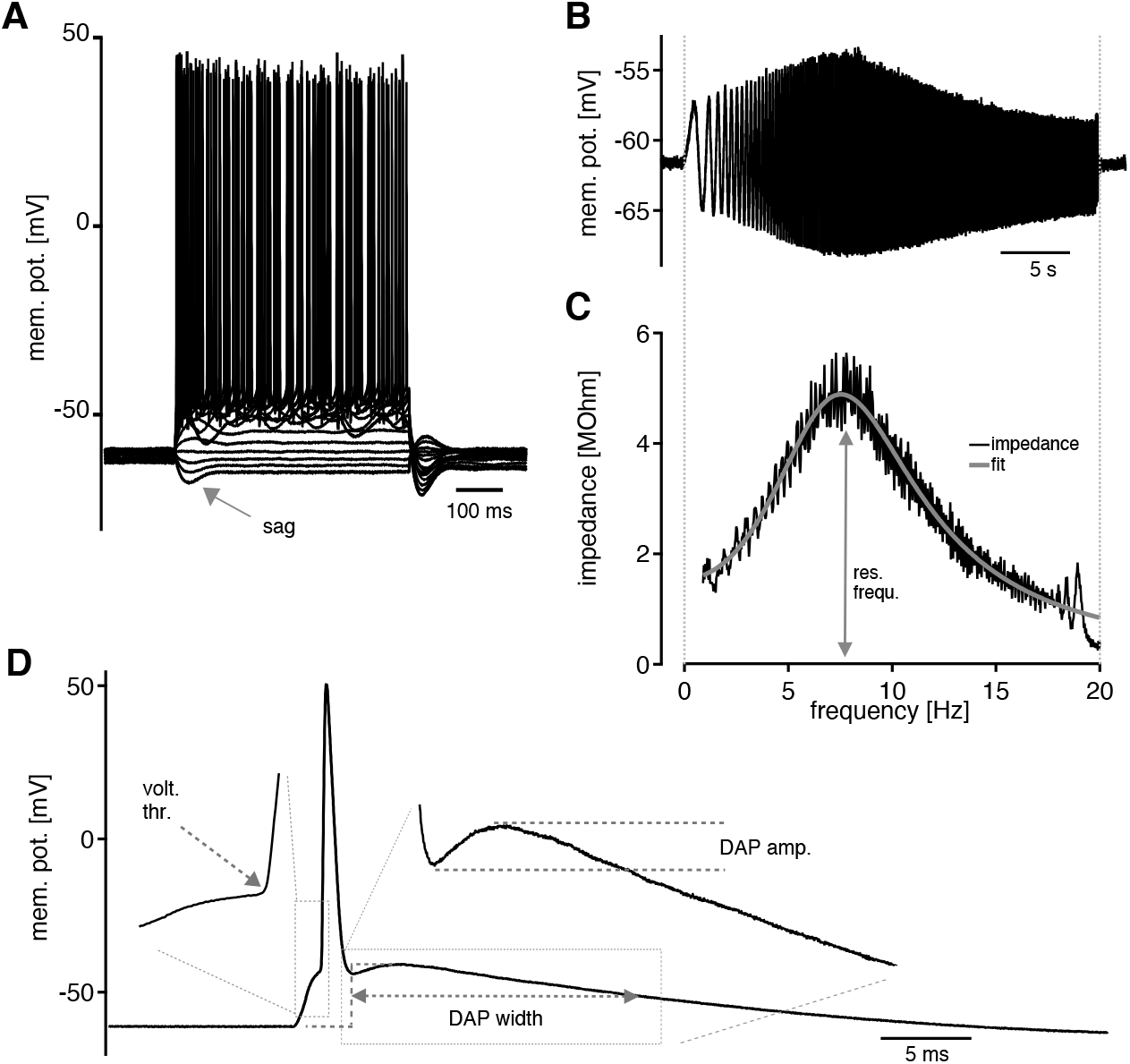
In-vitro electrical properties of mEC layer II cells in response to current-clamp stimulations using different standard protocols. ***A***, Voltage traces following step-current injections reveal sag potentials. The current injections ranged from -150 pA up to +800 pA and lasted 500 ms. ***B***, Membrane potential in response to a ZAP stimulation (0-20 Hz) lasting 30 seconds. ***C***, Estimation of the *resonance frequency* (*f*_*res*_), which was calculated from the impedance at different frequencies and measured as the maximum of the fitted curve (in grey). ***D*** Example voltage deflection in response to a short current injection that elicits a single AP. The left inset highlights the voltage threshold, right inset the *DAP amplitude*, defined as the voltage difference between the fAHP trough and the DAP peak. The *DAP width*, illustrated in the center, is defined as time between the fAHP trough and the point at which the DAP’s declining phase reaches the DAP half-maximum relative to the cell’s resting potential.

#### Latency and ISI1/ISI2 ratio

The latency of the first action potential following the onset of a step-current input was determined for inputs eliciting spike responses just above firing threshold. The same is true for measurements of the *ISI*1/*ISI*2 ratio, the quotient of the first and second inter-spike interval after stimulus onset.

#### Subthreshold resonance frequency

ZAPs (impedance amplitude profiles) are sinusoidally frequency-modulated current stimuli whose frequency *f*(*t*) increased linearly in time (Fig. 1*B*). They were used to estimate subthreshold resonance frequencies (Fig. 1*C*) and generated by the *Template Creator* which controls the Patchmaster by the “batch file control” protocol of *HEKA*.

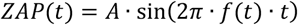

with

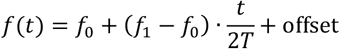

For the time-dependent frequency of the ZAP current, we used a range from 0-20 Hz (*f*_0_ = 0 Hz, *f*_1_= 20Hz) with a stimulus duration (*T*) of 30 seconds (as in Schreiber et al., 2004). This was done to ensure that the ZAP current changed slowly enough to obtain a precise estimation of the impedance. A current with a standard amplitude (*A*) of 0.1 nA was injected. By applying a constant offset current (offset) the cell’s membrane voltage could be adjusted to be either above or below spike threshold. The stimulation was repeated at least once. The recorded data were then transferred back from *Patchmaster* to the *Template Creator* and rapidly analyzed online to guide further experiments. This protocol was applied to all recorded cells as another standard protocol.

#### Voltage threshold and depolarizing afterpotential (DAP)

Single APs were elicited close to the AP threshold. To do so, a short depolarizing current consisting of a linearly rising ramp of duration 0.8 ms and a linearly descending ramp of 1.2 ms was injected, starting at the cell’s resting membrane potential. The current was increased in 100 pA increments until an action potential was elicited. This stimulus was used to measure a cell’s voltage threshold, defined as the inflection point of the voltage trace leading to an AP (see left inset in Fig. 1*D*). In addition, the DAP following the first supra-threshold response was analyzed. The *DAP amplitude* was determined as voltage difference between the fAHP trough and the following DAP peak (see right inset in Fig. 1*D*). The *DAP width* was calculated as the time between the fAHP trough and the point at which the declining phase of the DAP reaches the DAP’s half-maximum relative to the cell’s resting potential (see the horizontal arrow in Fig. 1*D*).

#### Analysis of electrical resonance

The analysis of electrical resonance responses to ZAP stimulations was performed as described in Erchova et al. (2004). First the frequency dependent impedance *Z*(*f*) was calculated as the ratio between the Fast Fourier Transform *V*(*f*) of the measured membrane potential *V*(*t*) divided by the Fast Fourier Transform *I*(*f*) of the injected ZAP current *I*(*t*),

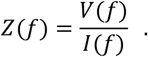

The impedance *Z*(*f*) is a complex variable and consists of a real and an imaginary part,

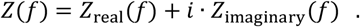

To estimate the resonance frequency the frequency-dependent amplitude |*Z*(*f*)| of the impedance,

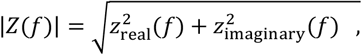

was fitted using the RCL model of Erchova et al. (2004) with four fit parameters (*a, b, c, d*),

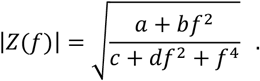

The resonance frequency (*f*_*res*_) estimated from this model corresponds to the frequency at which the maximum of the impedance curve is obtained (see Fig. *1D*),

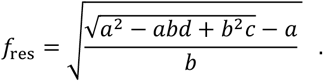

Frequencies of 1 Hz and less were excluded for the fits as estimating frequency-dependent impedances is less reliable in this regime (Erchova et al., 2004). For cells with low-pass properties *f*_*res*_ was set to 0 Hz. Another characteristic of resonators is the *Q* value, which captures the sharpness of the resonance. It is computed as the square-root of the ratio between the impedance’ size at the resonance frequency and its projected value at zero frequency,

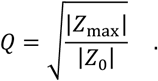

*Q* values above unity imply resonant behavior.

### Cluster analysis

To identify subpopulations amongst the recorded cells we carried out a quantitative cluster analysis. To obtain the optimal number of features which should be used for the clustering, we used a cross-validated factor analysis for dimensionality reduction and feature extraction. The performance of the factor analysis was evaluated by five-fold cross-validation and resulted in the highest average likelihood scores for seven features. Based on this analysis, we selected seven electrophysiological measures which represent distinct membrane properties and explain a large fraction (∼80%) of the observed variance in the data. On these selected features k-means clustering (Steinhaus, 1956) was performed for randomly chosen 50% of the data for faster convergence, a procedure that was repeated for 50 different random initial conditions. For each such clustering a *silhouette score* (Rousseeuw, 1987) was computed. The silhouette score measures the similarity of a cell’s data to the data points of the other cells within its own cluster and across clusters. For a given cell *i*, this score is calculated as

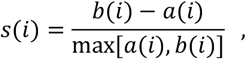

where *a*(*i*) is the average distance between the data point of cell *i* and all other data points in the cluster containing cell *i* and *b*(*i*) is the mean distance between the data of cell *i* and the data of all cells that are in the cluster nearest to the cluster cell *i* is residing in. The final score is the average across all cells and ranges from −1 to 1, with a high score indicating a high similarity within the cluster and low similarity across clusters.

To measure the clustering robustness, we compared the results from multiple clusterings. To this end, we clustered 50 random samples, each containing 50% of the data. Pairwise similarities between the clusterings were computed using the adjusted Rand (Rand, 1971) index as proposed by Vinh et al. (2009). For each of the resulting 1225 pairs, only data points that occurred in both samples were used. Finally, a *robustness score* was calculated as median over the values of the adjusted Rand index.

In addition to the k-means clustering a hierarchical Ward clustering was used to cross-check the clustering (Ward, 1963). Ward clustering is an agglomerative hierarchical clustering method, which means that each cell is first treated as a single cluster at the lowest hierarchical level. Clusters are then combined to form new clusters. This procedure is repeated recursively until all clusters are merged so that the relation between subclusters and superclusters can be analyzed in a quantitative manner. Whether clusters are merged or not is decided by an objective function, in our case the minimal variance within clusters. This approach is called Ward method. For both clustering procedures z-score normalized values (based on mean and standard deviation of each feature) were used to avoid potential biases due to variance differences between the different parameter axes.

To check the consistency of the cluster assignments across the two clustering algorithms we computed the robustness score as described above but this time measuring the 1225 similarities between Ward clusterings and k-means clusterings, separately for N=2 to N=6 clusters. By comparing similarity and robustness scores and assessing the most consistent cluster assignments across Ward- and k-means clustering an optimal number of clusters was estimated (N = 4). With this optimal cluster number the k-means algorithm was repeated on all data to get a cluster assignment for each cell.

### Data analysis and statistics

Data were processed and analyzed in *IGOR Pro* (WaveMetrics, Lake Oswego, OR), *Matlab* (MathWorks Inc., Natick, MA, USA) and *mecPhysio* (custom-made, J Nagele, LMU Munich). Statistics were calculated with *Prism 7* (Graphpad Software Inc., La Jolla, CA, USA). Group data are presented as mean ± standard deviation. Data were screened for normality with the Kolmogorov-Smirnov test and were then tested either by the parametric paired/unpaired t-test or by the non-parametric Mann-Whitney U test.

## Results

The morphology of principal cells in Layer II (LII) of the rodent medial entorhinal cortex (mEC) spans a wide range, from stellate cells (SCs) to pyramidal cells (PCs). Detailed analyses suggest two (mice, Fuchs et al., 2016), zero (mice, Winterer et al., 2017), or three (rats, Canto and Witter, 2012) intermediate cell classes. In mice, the electrophysiological properties of SCs, intermediate SCs (IMSCc) and intermediate PCs (IMPCs) vary gradually, whereas sag potentials and depolarizing afterpotentials (DAPs) in mice PCs seem to be far smaller than in the other cell types or in rat PCs (Fuchs et al., 2016). To better understand these results, we studied the electrophysiological properties of 531 rat mEC layer II neurons *in vitro*. Systematic cluster analyses helped us to identify and characterize distinct subpopulations of LII mEC cells and allowed us to estimate the variability of DAP parameters within and between the clusters. The findings provide a comprehensive picture of the physiology of rodent LII mEC cells that is consistent with *in vivo* data (Domnisoru et al., 2013; Latuske et al., 2015; Csordás et al., 2020).

### Electrophysiological characterization

Whole-cell current-clamp recordings were performed to characterize the sub- and supra-threshold properties of mEC LII neurons, as described previously (Alonso and Klink, 1993; Klink and Alonso, 1997; van der Linden and da Silva, 1998; Canto and Witter, 2012, Alessi et al., 2016). The resting membrane potential *E*_*rest*_ was estimated by averaging the membrane potential over 200 ms in the absence of any injected current (see Methods). IV protocols were used to identify sag potentials in response to a hyperpolarizing current injection of -150 pA. The sag was calculated as the voltage difference between the mean membrane potential at steady state (last 200 ms) and the cell’s maximum hyperpolarization (Fig. 1*A*). A stepper protocol was applied to measure the membrane resistance *R*_*in*_ and the membrane time constant *τ*. As detailed in the Methods, we used a ZAP (impedance amplitude profile) stimulus to determine whether the membrane potential displayed an electrical resonance (Fig. 1*B*). The frequency at which the fitted impedance curve has its maximum was labeled as *f*_*res*_ (Fig. 1C). Finally, the cell’s voltage threshold and depolarizing afterpotential (DAP) (Fig. 1*D*) were extracted from the voltage response to a short current ramp injection. The DAP occurs between a fast afterhyperpolarisation (fAHP) and a medium-range afterhyperpolarization (mAHP), the slow recovery phase of the membrane potential toward the resting potential, as illustrated in Fig. 1*D*.

Our first goal was to examine the properties of DAPs in the different mEC layer II cell types. Therefore, single APs were elicited close to the threshold (see Methods). From these recordings two DAP characteristics were defined: *DAP amplitude* and *DAP width*. The *DAP amplitude* was calculated as the voltage difference between the local minimum of the fAHP and the following depolarizing peak, as shown by the arrow in the magnified inset in Fig. 1*D*. The *DAP width* was computed as the time interval between the fAHP trough (see dotted arrow in Fig. 1*D*) and the point at which the declining phase of the DAP reaches half-maximum relative to the cell’s resting potential, as indicated by the horizontal arrow in Fig. 1*D*.

Correlations between DAP amplitude or DAP width and basic electrophysiological parameters were calculated in terms of Spearman correlations. The DAP amplitude correlated positively with the time interval between the AP and DAP maxima (r=0.60, p<0.0001): in cells with larger DAPs the DAP maximum occurs at later times. The DAP width also correlates positively with the time interval between AP and DAP maximum (r=0.60, p<0.0001) and negatively with the resonance frequency (r=-0.37, p<0.0001). The last two results are consistent with the view that there is a gradient between “slower” and “faster” neurons regarding all three temporal cell characteristics.

### Cluster analysis

We next investigated whether the recorded neurons belonged to distinct subpopulations. To this end, we performed a quantitative cluster analysis (see Methods) based on physiological parameters characterizing sub- and supra-threshold responses. On purpose, we did *not* include DAP-related parameters so as to be able to test whether those parameters can be inferred from the other data. As a cross-validated factor analysis (see Methods) suggested seven parameters, we chose seven distinct electrophysiological quantities that explain a large fraction (∼80%) of the observed variance in the data, namely the membrane resistance *R* and time constant *τ*, resonance frequency *f*_*res*_, *Q-*value, first-AP latency at firing threshold, *ISI*1/*ISI*2 ratio, and AP width.

For these parameters, k-means clustering revealed the silhouette and robustness scores listed in Table 1. The silhouette score measures how similar the data points are within a cluster compared to the similarity to other clusters. This score ranges between −1 and 1. A second score, the robustness score, was used to assess the clustering robustness: A robustness score of zero corresponds to non-robust cluster assignments that fluctuate randomly across repeated clusterings on 50%-subsets of the data. A score of one means that the cluster assignments are always equal across repetitions. To scrutinize these results we repeated the cluster analysis using an alternative approach, hierarchical Ward clustering (Methods).

**Table 1.** Silhouette and robustness scores of k-means and Ward clusters for different cluster numbers, together with robustness scores that test the similarity of clusters from either method.

|  | Number N of clusters |  |  |  |  |
| --- | --- | --- | --- | --- | --- |
|  | N = 2 | N = 3 | N = 4 | N = 5 | N = 6 |
| **k-means clustering** |  |  |  |  |  |
| Silhouette score | 0.404 | 0.327 | 0.315 | 0.253 | 0.222 |
| Robustness score | 0.956 | 0.954 | 0.813 | 0.638 | 0.571 |
| **Hierarchical Ward clustering** |  |  |  |  |  |
| Silhouette score | 0.535 | 0.397 | 0.353 | 0.263 | 0.229 |
| Robustness score | 0.596 | 0.724 | 0.719 | 0.544 | 0.499 |
| **Comparison: k-means vs. Ward clustering** |  |  |  |  |  |
| Robustness score | 0.614 | 0.785 | 0.798 | 0.375 | 0.378 |

On a coarse level, this clustering (see Fig. 2) reveals two main branches, corresponding to cells with “stellate” and “pyramidal” characteristics, respectively. On a finer scale, first the stellate branch separates into C1 (SC) and C2 (IMSC). On a yet finer scale, marked by the dashed line in Fig. 2, the pyramidal branch splits into C3 (IMPC) and C4 (PC). The late separation underscores the higher similarity within the pyramidal branch compared to the stellate branch.

**Figure 2.**
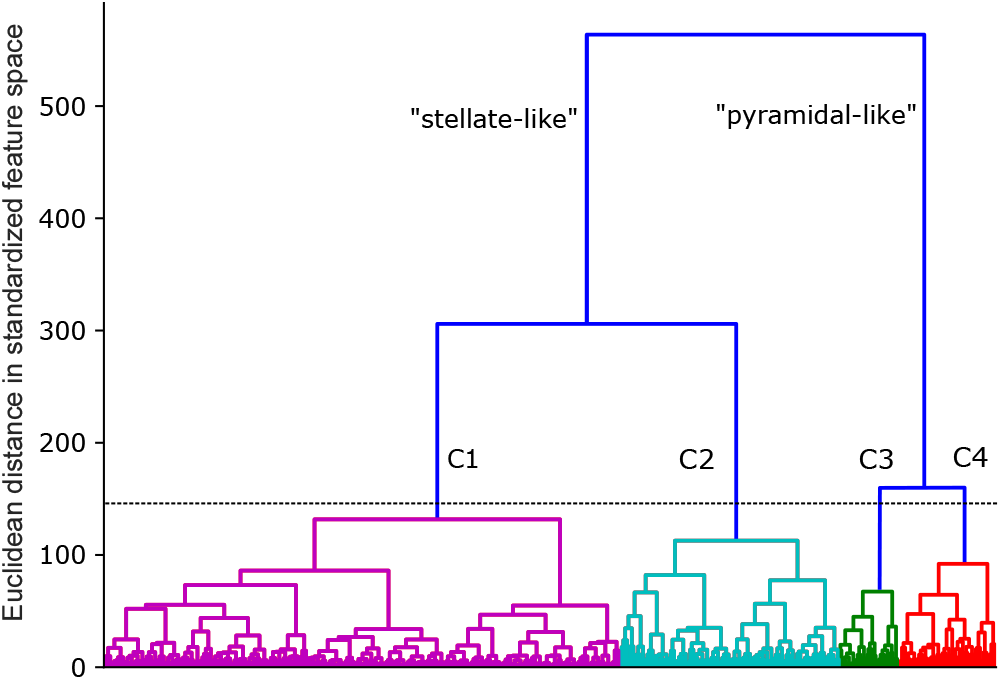
Dendrogram illustrating the arrangement of the clusters computed by hierarchical Ward clustering. The dashed line illustrates the appearance of four clusters, which are distinctly colored.

As expected, the two cluster algorithms resulted in different cluster scores (Table 1). However, their overall dependency on the cluster numbers is rather similar. For both, k-means clustering and hierarchical Ward clustering, the silhouette and robustness scores drop strongly once more than four clusters are considered, and so does the similarity between the clusters obtained with both algorithms. Thus N=4 seems to be an upper limit for the number of distinct cell clusters. On the other hand, clusterings with N=2 result in particularly high values for the two silhouette scores. However, the robustness of the comparison between k-means and Ward clustering is much lower for N=2 than for N=3 or N=4. This implies that clusters obtained from either algorithm are more reliable for N=3 or N=4, compared to N=2. In addition, larger cluster numbers should be preferred because the data in this study were obtained from multiple animals, were not controlled regarding the dorsoventral gradient within the mEC and may also differ in the state of the respective brain slices. Such unaccounted variabilities in the cells’ properties tend to blur cluster boundaries and result in an underestimated cluster number. In what follows, we therefore focus on N=3 and N=4.

To decide whether three or four clusters capture the data better one may inspect the features on which the clustering is based on. This is shown in Fig. 3, which plots the cumulative histograms of the seven selected features for three (N=3) and four (N=4) clusters using the k-means algorithm. Cluster C1 of the three-cluster solution is highly reminiscent of cluster C1 of the arrangement with four clusters. Cluster C2 of the three-cluster solution overlaps with cluster C2 of the four-cluster situation. Cluster C3 of the partition in three clusters is very similar to clusters C3 and C4 of the four-cluster soluion, suggesting that three clusters suffice to describe the variability in most electrophysiological data. But comparing membrane time constants (Fig. 3*C* vs. Fig. 3*D*) and first latencies at firing threshold (Fig. 3*I* vs. Fig. 3*J*) indicates that clusters C3 and C4 of the four-cluster arrangement differ and can be separated. A partition into four clusters is also supported by the observation that the similarity between k-means and Ward clustering peaks at N=4 (Table 1).

**Figure 3.**
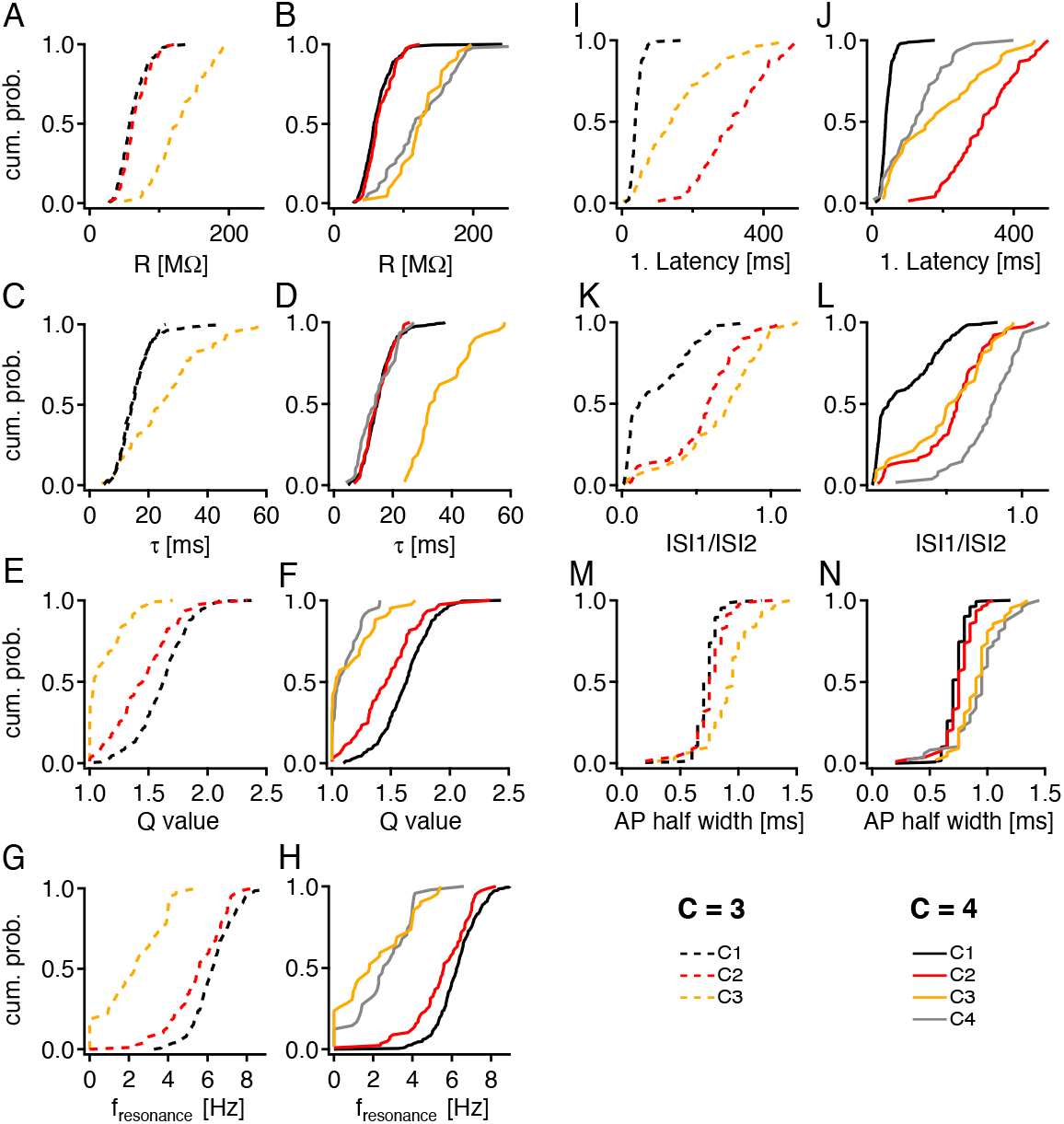
Comparison between three (N=3) and four clusters (N=4) generated by k-means clustering. ***A*** and ***B***, membrane resistance *R*; ***C*** and ***D***, membrane time constant *τ*; ***E*** and ***F***, Q value; ***G*** and ***H***, resonance frequency *f*_*res*_; ***I*** and ***J***, first-AP latency at firing threshold; ***K*** and ***L***, ISI1/ISI2 ratio; ***M*** and ***N***, AP half width. Colored lines indicate the different clusters.

Analyzing the cells’ allocation to the four K-means clusters and four Ward clusters (see Table 2) shows that both clusterings agree with high accuracy regarding the stellate versus pyramidal dichotomy (32 false assignments compared to 385 correct ones). Within the stellate clusters (C1 and C2), both clusterings coincide in 308 cases and differ in 12 cases only, in agreement with the early split between C1 and C2 for the hierarchical Ward clustering. On the other hand, there is a high level of mixing when it comes to the C3 versus C4 clusterings (35 correct, 30 incorrect), as might be expected from the late splitting of the C3 and C4 Ward clusters.

**Table 2.** Comparison between k-means clustering and hierarchical Ward clustering. There is high agreement between the classifications proposed by the two different algorithms for Clusters C1 and C2 but strong mixing between Clusters C3 and C4.

| Cluster | Ward C1 | Ward C2 | Ward C3 | Ward C4 |  |
| --- | --- | --- | --- | --- | --- |
| K-means C1 | 240 | 11 | 0 | 1 | $\Sigma = 252$ |
| K-means C2 | 1 | 68 | 0 | 7 | $\Sigma = 76$ |
| K-means C3 | 0 | 6 | 17 | 19 | $\Sigma = 42$ |
| K-means C4 | 0 | 18 | 11 | 18 | $\Sigma = 47$ |
| | $\Sigma = 241$ | $\Sigma = 103$ | $\Sigma = 28$ | $\Sigma = 45$ | $\Sigma = 417$ |

We conclude that the measured data are best explained by assuming four cell clusters. We should keep in mind, however, that apart from differences in first-spike-latency behavior, cells in clusters C3 and C4 possess similar electrophysiological characteristics, as also reflected in the strongly reduced separability of these two clusters in the comparison between k-means vs. Ward clustering.

### Electrophysiological characteristics of the cell clusters

To compare the selected features and the electrophysiological properties of the four clusters (see Table 3) a non-parametric Kruskal-Wallis test was used. All tested parameters are significantly different when the four clusters are compared (Voltage threshold: p=0.0003; all other parameters: p<0.0001). The cumulative histograms in Fig. 4 and the data listed in Table 3 clearly illustrate that cells in clusters C1 and C2 are similar in their resting membrane potential (Fig. 4*A*), voltage threshold (Fig. 4*B*), capacitance (Fig. 4*C*), and sag potential (Fig. 4*D*). Clusters C3 and C4 are also very similar in their basic electrophysiological properties (see Table 3).

**Table 3.** Basic electrophysiological characteristics of the four clusters. Values are means ± SD. The numbers differ significantly from cluster to cluster (Spike threshold: p=0.0003; all other parameters: p<0.0001) as calculated with a non-parametric Kruskal-Wallis test. The numerical values in the six lower rows differ from those in Fuchs et al. (2016). However, the inter-cluster trends are similar.

| Electrophysiological measure | C1 (SC) | C2 (IMSC) | C3 (IMPC) | C4 (PC) |
| --- | --- | --- | --- | --- |
| Membrane time constant $\tau$ (ms) | $15.2 \pm 5.4$ | $15.0 \pm 4.6$ | $36.6 \pm 9.4$ | $15.0 \pm 7.2$ |
| Membrane resistance $R$ (MΩ) | $62.27 \pm 22.5$ | $66.13 \pm 19.3$ | $126.1 \pm 34.0$ | $126.3 \pm 54.0$ |
| Resonance frequency $f_{res}$ (Hz) | $6.25 \pm 1.13$ | $5.51 \pm 1.47$ | $2.15 \pm 1.81$ | $2.57 \pm 1.49$ |
| Q value | $1.62 \pm 0.21$ | $1.47 \pm 0.27$ | $1.16 \pm 0.20$ | $1.1 \pm 0.12$ |
| 1st AP latency (ms) | $41.7 \pm 19.0$ | $318.6 \pm 93.6$ | $190.0 \pm 131.0$ | $127.6 \pm 83.3$ |
| ISI1/ISI2 ratio | $0.22 \pm 0.2$ | $0.55 \pm 0.24$ | $0.52 \pm 0.27$ | $0.79 \pm 0.20$ |
| AP half width (ms) | $0.77 \pm 0.01$ | $0.75 \pm 0.14$ | $0.92 \pm 0.17$ | $0.9 \pm 0.24$ |
| Resting potential $E_{rest}$ (mV) | $-62.8 \pm 4.1$ | $-64.3 \pm 4.5$ | $-69.1 \pm 8.6$ | $-69.9 \pm 7.7$ |
| Sag amplitude (mV) | $2.38 \pm 0.85$ | $2.41 \pm 0.98$ | $1.89 \pm 1.56$ | $1.62 \pm 2.01$ |
| Spike threshold (mV) | $-41.62 \pm 5.95$ | $-40.41 \pm 6.68$ | $-37.33 \pm 11.4$ | $-37.07 \pm 8.02$ |

**Figure 4.**
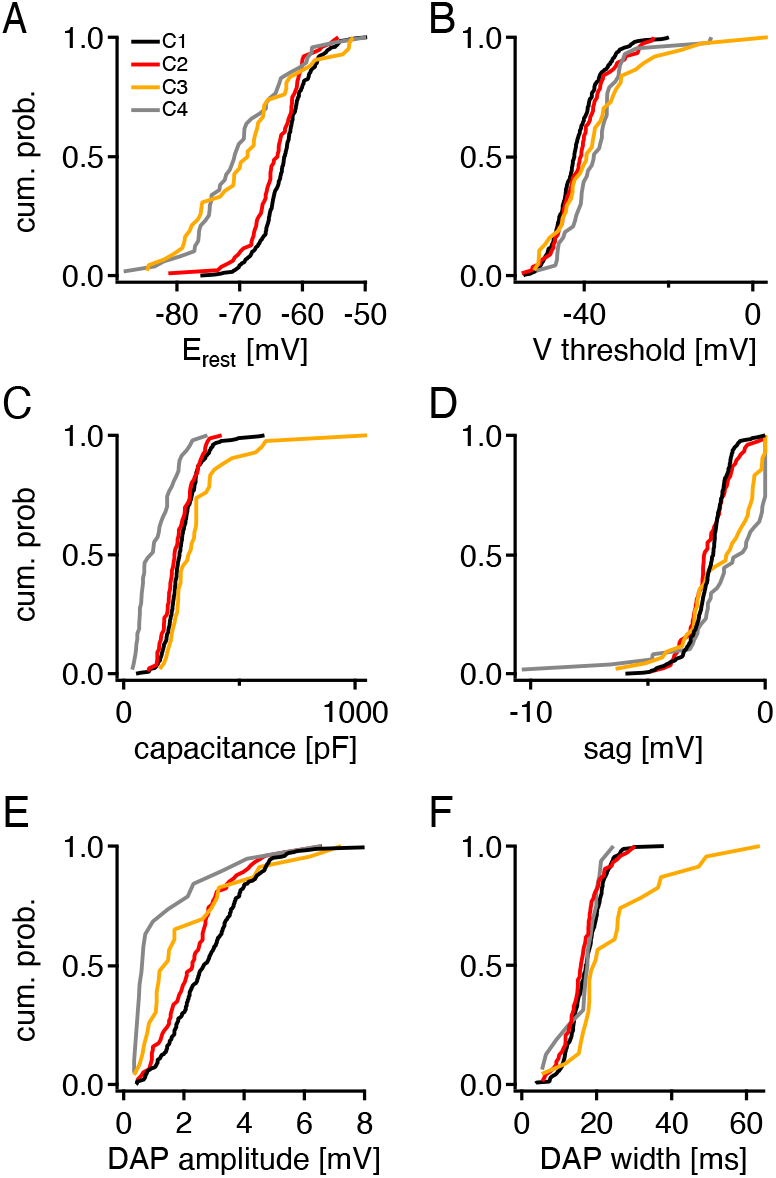
Electrophysiological properties within each of the four k-means clusters. Colored lines indicate the different clusters.

A key parameter, which distinctly separates the four clusters, is the latency of the first suprathreshold response shown in Fig. 3*J*. The majority of C1 cells have a very short latency, whereas cells in the C2 cluster have a very long latency, even longer than that observed for C3 and C4. On the other hand, there is quite a substantial overlap (Fig. 4*D*) in the case of the sag, which is highly similar for about half of the cells in each cluster. For the other half, the sag is either mostly in the 1.5-2 mV range (C1 and C2) or below 1.5 mV (C3 and C4).

These results suggest that C1 cells are SCs, which show a pronounced sag, resonate at frequencies in the 5-9Hz theta range and have a very short latency to the first spike at firing threshold, a consequence of the sag response to depolarizing currents (Alonso and Klink, 1993). In contrast, cells in cluster C4 are reminiscent of PCs as they have larger membrane resistances and act as a low-pass filter (Alonso and Klink, 1993; van der Linden and da Silva, 1998; Erchova et al., 2004; Canto and Witter, 2012). Around half of the C4 cells show only a very small or no sag response and the latency to the first spike is much longer than that of C1 cells, again pointing to PCs. Cells in Cluster C2 are stellate-like but with long latencies, corresponding to the intermediate SCs reported by Fuchs et al. (2016).

### Depolarizing afterpotentials

A key motivation for this study was the question whether the amplitude and width of depolarizing afterpotentials vary significantly between the different mEC LII cell clusters. To answer this question in an unbiased manner, DAP features were excluded for the clustering procedure.

A comparison of DAP amplitudes and widths across all four clusters shows significant differences, based on a non-parametric Kruskal-Wallis test (Table 4). The cumulative histograms of Fig. 4*E* illustrate that the majority of cells in clusters C1 and C2 exhibit a pronounced DAP (> 2 mV). C3 cells display a somewhat smaller DAP, whereas more than 40% of the C4 cells show either a very small DAP (< 0.5 mV) or no DAP at all; only a small fraction of C4 cells (∼20 %) elicit pronounced DAPs above 2 mV (Fig. 4*E*). An inspection of the DAP width shows a less clear separation. Only cluster C3 deviates from the other clusters in that it contains a substantial fraction of cells with wide DAPs (see Fig. 4*F*). Note, however, the large variability (SD = 13.6 ms), potentially due to outliers in the small sample.

**Table 4.** Means ± SD of DAP amplitudes and widths for the four cell clusters. Differences are significant based on the non-parametric Kruskal-Wallis test.

|  | Cluster |  |  |  |  |  |  |  | p value |
| --- | --- | --- | --- | --- | --- | --- | --- | --- | --- |
|  | C1 (SC) |  | C2 (IMSC) |  | C3 (IMPC) |  | C4 (PC) |  |  |
| DAP Amplitude (mV) | 2.61 | $\pm 1.5$ | 2.37 | $\pm 1.3$ | 2.2 | $\pm 1.85$ | 1.41 | $\pm 1.6$ | $< 0.0001$ |
| DAP Width (ms) | 17.4 | $\pm 6.2$ | 16.1 | $\pm 5.2$ | 25.1 | $\pm 13.6$ | 16.3 | $\pm 5.4$ | 0.0042 |

Together, these findings support the view that DAPs in layer-II of mEC are not restricted to certain cell groups but rather form a broad continuum across all principal neuron types.

## Discussion

Slice experiments have shown that depolarizing afterpotentials (DAPs) arise in a majority of principal cells in mEC Layer-II (Alonso and Klink, 1993; Canto and Witter, 2012, Fuchs et al., 2016, Winterer et al., 2017). DAPs also shape the cells’ burst behavior: during DAPs the AP current threshold is strongly reduced such that the average excitability increases by over 40% (Alessi et al., 2016). Whole-cell recordings from mice moving on a virtual linear track (Domnisoru et al., 2013) show that *in vivo*, too, DAPs play a decisive role for burst firing in mEC Layer-II neurons: Cells with DAPs are bursty and their intra-burst ISIs are compatible with the DAP mechanism (Csordás et al., 2020). These observations trigger the question whether DAPs are exhibited by all cell types in mEC Layer-II or whether there are certain cell types that do only show small DAPs or no DAPs at all, as suggested by Fuchs et al. (2016) and Winterer et al. (2017).

To answer this question, one needs to first classify the different cell types in an unbiased manner. We did so by using k-means clustering and, as a control, hierarchical Ward clustering. As suggested by a cross-validated factor analysis, the clusterings were based on seven parameters, chosen as membrane resistance *R*, membrane time constant *τ*, resonance frequency *f*_*res*_, *Q*-value, first-AP latency at firing threshold, *ISI*1/*ISI*2 ratio, and AP width. To minimize any potential bias, we did *not* include DAP properties within this group of electrophysiological variables used for the clusterings.

The cells analyzed in this study originated from multiple animals, were not controlled regarding their dorsoventral location within the mEC and may differ in the vital state of the respective brain slices. All these factors introduce additional variability in the cells’ response properties that tends to blur cluster boundaries and to reduce the number of clusters. However, the success of the cluster analysis suggests that although parameters within each cell class vary measurably, as has been shown for stellate cells (Giocomo and Hasselmo, 2008; Pastoll et al., 2020), taken together they nevertheless give rise to a pronounced cluster structure. In fact, the cluster analysis suggests four clusters that turn out to be reminiscent of the four clusters of Fuchs et al. (2006) for mice LII mEC principal neurons. There is, however, one striking difference: In mice, DAPs of PCs are almost non existing (0.04 ± 0.05 mV, mean ± SD) according to Fuchs et al. (2016) or rather small (0.26 ± 0.44 mV, mean ± SD) as in Winterer et al. (2017). In contrast, our data from rats suggest much larger values (1.41 ± 1.6 mV, mean ± SD), in agreement with the findings by Canto and Witter (2012) for weak stimulation, as shown in Table 5.

**Table 5.** Comparison of DAP amplitudes from different studies. Unless otherwise noted, DAP amplitudes are defined as difference between DAP peak and fAHP trough. The results of Alessi et al. (2016) include data about the DAP dependence on the holding potential V_h_ prior to triggering the DAP, Canto and Witter (2012) provide information about the dependence on the injected current triggering the DAP. To facilitate the comparison between the different studies, for the data from Canto and Witter, means and SDs are calculated by taking the entire cell populations into account, not only the cells with non-zero DAP amplitudes, as was done by those authors. The standard deviations of the data from Fuchs et al. (2016) are calculated from the published standard errors of the means. Data in Alessi et al. (2016): visual inspection of their Fig. 2; data in Winterer et al. (2017): private communication.

| DAP amplitude (mV)<br>• all data: mean ± SD | SC | IMSC | IMPC | PC |
| --- | --- | --- | --- | --- |
| Rat (this paper)<br>• $V_h = V_{rest}$ | $2.61 \pm 1.5$ | $2.37 \pm 1.3$ | $2.2 \pm 1.85$ | $1.41 \pm 1.6$ |
| Rat (Alonso & Klink, 1993)<br>• $V_h$ not reported | $1.1 \pm 1.0$ | — | — | $0.4 \pm 0.6$ |
| Rat (Canto & Witter, 2012)<br>• $V_h$ not reported<br>• DAP amplitude measured relative to $V_{thresh}$ | $2.80 \pm 1.21$ for weak stimulation<br>$3.61 \pm 1.60$ at +200pA | — | — | $1.49 \pm 1.01$ for weak stimulation<br>$2.79 \pm 1.60$ at +200pA |
| Rat (Alessi et al., 2016)<br>• two $V_h$ values explored | 3.3 at $V_h = -67$mV<br>6.0 at $V_h = -82$mV | — | — | 0.1 at $V_h = -87$mV<br>1.8 at $V_h = -67$mV |
| Mouse (Fuchs et al., 2016)<br>• $V_h$ not reported | $2.35 \pm 0.64$ | $1.46 \pm 0.56$ | $1.56 \pm 0.30$ | $0.04 \pm 0.05$ |
| Mouse (Winterer et al, 2017)<br>• $V_h$ not reported | $1.97 \pm 1.82$ | — | — | $0.26 \pm 0.44$ |

In principle, the discrepancy between the PC data from mice and rats could reflect a species difference. The roughly six-fold deviation between the DAP amplitudes in mouse PCs reported by Fuchs et al. (2016) and those by Winterer et al. (2017) does, however, suggest that the DAP amplitudes in PCs depend sensitively on experimental details or exhibit a pronounced cell-type intrinsic variability. This conjecture is supported by the observation that there is also a more than three-fold difference between the DAP amplitudes of rat PCs in Alonso and Klink (1993) compared with those in Canto and Witter (2012) or the present study. Similarly, a large fraction of PCs do not show any measurable DAPs: 45% in Alonso and Klink (1993), roughly 25% in Canto and Witter (2012), much higher percentages than those for stellate cells: 14% in Alonso and Klink, and 15% in Canto and Witter. With more than 40% of the pyramidal DAP amplitudes below 0.5 mV, despite a mean amplitude of more than 1.4 mV, our data points to a large variability of the pyramidal-cell DAPs, too, as do the large standard deviations in the other studies. The scatter plots of the PC data in Fuchs et al. (2016) and Winterer et al. (2017) might even suggest a binary separation of (almost)-zero and clearly non-zero DAP amplitudes.

There is yet another possible explanation for the discrepancy between our results and those of Fuchs et al. (2016) or Winterer et al. (2017). Fuchs et al. as well as Winterer et al. based their cell classification on a principal component analysis that included the DAP amplitude as one parameter. Depending on the distribution of DAP amplitudes, this might have led to a pronounced axis along the DAP-amplitude direction so that all cells with small-amplitude DAPs are grouped together, irrespective of their overall location in parameter space. It was also for this reason that, on purpose, we did *not* include any DAP properties in the clustering procedure.

In conclusion, our findings indicate that DAPs are not specific to a certain type of principal cells in layer II of medial entorhinal cortex but rather distributed across all principal cell types. Following an action potential triggered at rest, mean DAP amplitudes range from about 1.4 mV in pyramidal cells, to 2.2 mV in intermediate pyramidal cells, to 2.4 mV in intermediate stellate cells, to about 2.6 mV in stellate cells. All these mean values come with noticeable standard deviations of around 1.5 mV, underscoring a pronounced variability around the pyramidal-to-stellate axis – a variability that is particularly strong for pyramidal cells. The overall cell-to-cell variability is also observed in *in-vivo* data from mice running in virtual linear tracks (Domnisoru et al., 2013) or real two-dimensional arenas (Latuske et al.,2015) and suggests that DAP phenomena depend sensitively on the cell’s physiological state (Csordás et al., 2020) and can be modulated to shift the cell between regular firing and a DAP-induced burst mode.

## Acknowledgements

This work has been supported by the German Research Foundation (via CRC 870). We thank D Csordás, C Fischer and M Stemmler for stimulating discussions and J Winterer for sharing unpublished data.

